# Initial identification of genomic islands in *Taylorella equigenitalis* and *Taylorella asinigenitalis* and their distribution in isolates from around the world

**DOI:** 10.1101/2025.11.26.686905

**Authors:** Jessica Hicks, Ganwu Li, Sandrine Petry, Fabien Duquesne, Jun-Gu Choi, Gundrun Overesch, Kristina Lantz, Suelee Robbe-Austerman, Xiaoqiu Huang

## Abstract

*Taylorella equigenitalis*, the causative agent of contagious equine metritis (CEM), is endemic in several countries throughout the world and impacts the global movement of horses. It presents with symptoms ranging from non-clinical to endometritis, vaginitis, and acute infertility in females while males remain asymptomatic. Studies of the disease have focused primarily on understanding pathology, transmission dynamics, diagnostics, and phenotypic and genotypic characteristics. The goal of this study was to establish a more comprehensive global phylogenetic structure and generate reference-quality genomes for each major clade of the tree where such data were absent. Seven of these genomes along with 44 other publicly available genome assemblies, including *Taylorella asinigenitalis*, were then used to predict genomic islands. Exploration of the NCBI annotations in the genomic islands indicate links to virulence and ecological fitness. An additional 97 *T. equigenitalis* and 48 *T. asinigenitalis* short-read isolates from this study and 215 publicly available genomes were used to examine distribution of each genomic island across the phylogenetic tree for evidence of horizontal movement. The stability of these genomic islands in their natural environment was evaluated utilizing 141 isolates recovered from a recent CEM outbreak in the United States. The results obtained in this study demonstrate mobile genomic islands in *T. equigenitalis* genomes, sites of concern for their contribution to the potential virulence of the organism.

## 1 Introduction

Contagious equine metritis (CEM), a disease caused by the bacterium *Taylorella equigenitalis*, was first reported in 1977 and remains an international trade concern in the horse industry [1]. In females, the disease manifests as copious vaginal discharge, vaginitis, and endometritis, sometimes resulting in acute infertility; but symptoms present with varying degrees of clinical illness from severe to subclinical. Males are asymptomatic carriers of the bacteria making them a significant concern in disease transmission [2–5]. Global movement of horses and the often-low numbers of colonized bacteria on males allows many opportunities for the spread of *T. equigenitalis*.

Several studies have sought to characterize *T. equigenitalis* using various approaches, ranging from in vitro cellular infiltration to genotyping techniques such as pulse-field gel electrophoresis (PFGE) and multi-locus sequence typing (MLST) [6–8]. These methods have been instrumental in grouping strains and improving epidemiological tracing but fall short in molecular characterization of individual strains and identifying their potential virulence factors. A recent phylogenetic analysis of 28 *T. equigenitalis* genomes revealed that, on average, 6.6% of each genome potentially contained recombination events [9]. Genomic comparisons between *T. equigenitalis* isolates have uncovered differences among individual genomes, highlighting discrete regions of diversity and potential mobile elements [10–12]. Some genes within these variable regions are associated with virulence, including hemagglutinin proteins, which function as mammalian cell adhesins in many bacteria; type IV secretion systems (T4SSs), which mediate effector protein delivery across the cell membrane and facilitate the uptake of environmental proteins or DNA; and CRISPR-Cas systems [10–14]. However, it remains unclear whether these variable regions arise from mobile elements or other sources of genetic variation. Moreover, these genomic comparisons have been limited to a small number of isolates, potentially under-representing the species’ genomic diversity. A comprehensive study analyzing isolates from diverse geographic regions is yet to be conducted. Expanding our understanding of *T. equigenitalis* genomics, virulence factors, and their mobility between strains could lead to advancements in diagnostic assays and a deeper comprehension of the organism’s disease cycle.

*Taylorella asinigenitalis*, the only other species in the genus Taylorella, is generally considered non-pathogenic [15–17]. However, some strains have shown the potential to cause mild clinical illness and infection of mares [18, 19]. Relatively little has been done to establish genomic diversity or understand phylogenetic structure, and relatively little public data exists for this organism. An initial genome comparison of a single strain of *T. asinigenitalis* (MCE3) with a single strain of *T. equigenitalis* (MCE9) indicated a large overlap in genes, with 1322 of the 1534 annotated genes in TA being shared with TE; but it is unknown if this overlap includes genetic mechanisms that contribute to virulence. Also, a large inversion in the *T. asinigenitalis* genome relative to *T. equigenitalis* was noted [11].

Genomic islands (GEIs) are broadly defined as large, mobile gene clusters within a genome and can include many types of mobile elements, e.g., prophages, integrative conjugative elements, integrons, conjugative transposons, among others [20, 21]. GEIs can considerably enhance the genetic potential of an organism and have several functional classifications such as pathogenicity islands (PAIs), ecological islands, and other types related to metabolism, symbiosis, and antimicrobial resistance [20, 22]. GEIs are considered to contribute to genomic plasticity with their ability to easily move sets of genes in and out of the genome as needed in order to adapt to environmental challenges [20]. Often, GEIs occur near tRNAs, differ in GC content from the rest of the genome, and contain mobility genes such as integrases or translocases. Prediction of GEIs is accomplished through a variety of different algorithms used to detect regions that meet these characteristics: comparison to known GEIs, presence of known mobility genes in combination with nucleotide biases, and hidden Markov model approaches [23, 24].

The first goal of this study was to establish a more comprehensive global phylogenetic structure. Seven new clades consisting of the sequences from this study were added to the *T. equigenitalis* tree and complete, reference-quality genomes for these clades were created. Secondly, the reference sequences along with 45 other publicly available genome assemblies, including *T. asinigenitalis*, were analyzed to predict GEIs and examine gene content. While NCBI annotations of the predicted GEIs reveal potential links to virulence and ecological fitness, the abundance of hypothetical proteins underscores the potential for additional, unidentified functions. To investigate conservation and distribution of the GEIs across the phylogenetic tree, an additional 97 *T. equigenitalis* and 48 *T. asinigenitalis* short-read isolates from this study alongside 215 publicly available genomes were examined. Thirdly, to explore this dynamic in natural settings, 141 *T. equigenitalis* isolates recovered from a recent CEM outbreak in the United States were evaluated for presence and absence of the GEIs, as well as the combinations of GEIs found in these isolates. The results obtained in this study demonstrate the mobility of the predicted GEIs in *T. equigenitalis* genomes and their contribution to the potential virulence of the organism.

## 2 Materials and Methods

### 2.1 Isolate Selection

To better characterize the global phylogeny, collaborators contributed isolates collected from Belgium, France, the Netherlands, South Africa, South Korea, Switzerland, the United Kingdom, and the USA. Each country selected isolates that represented the most diversity for their isolate collection, either by MLST or by geographical region, or sent their entire collection. In addition to a previously sequenced culture collection of *T. equigenitalis* isolates from domestic and imported detections in the USA, additional isolates more recently recovered from imported horses (n= 97 *T. equigenitalis*) and from a recent outbreak (n=141 *T. equigenitalis*) within domestic ponies and horses were selected. All *T. asinigenitalis* isolates from the NVSL reference culture collection and a single isolate submission from France were also included in this study (n= 48 *T. asinigenitalis*).

### 2.2 Culture, DNA Extraction, and Sequencing

Isolates were obtained from bacteriological culture at the National Veterinary Services Laboratories (NVSL) in Ames, Iowa, or shipped as frozen isolates. Isolates were subcultured on chocolate Eugon agar with 5% horse blood from frozen culture and incubated for 48-72 hours at 37*^◦^*C in 5% CO2. Colonies were selected and extracted using either the Epicenter Masterpure kit (Epicentre, Madison, WI) or a Promega Maxwell with Maxwell Whole Blood DNA Kit (Promega Corporation, Madison, WI). DNA library preparation was performed with a Nextera XT kit (Illumina, San Diego, CA), and sequencing was performed on an Illumina MiSeq with a 2x250bp chip or an Illumina NextSeq with a 2x150bp or 2x250bp chip. A minimum depth of 70X coverage was ensured. Isolates selected for long-read sequencing were propagated in the same manner, extracted using PureGene (Qiagen, Hilden, Germany), and sequenced on a PacBio Sequel (Pacific Biosciences, Menlo Park, CA, USA).

### 2.3 Evaluation, Assembly, and Phylogenetic Analysis

Short-read sequences were screened for purity using Kraken and a custom database that includes the NCBI RefSeq database as well as potential host references [25]. Short-read sequences were then de novo assembled with Spades [26]. Additional *T. equigenitalis* and *T. asinigenitalis* sequences in NCBI were downloaded and with the Spades assemblies of both *Taylorella* species, a reference independent maximum likelihood phylogenetic tree was generated using kSNP with a 0.8 majority setting [27]. This analysis was used to select isolates for long-read sequencing to develop references for each major branch of the tree. Once this sequencing was completed, a TE-only tree was also constructed using kSNP.

Long-read sequences were assembled with Canu and Unicycler [28, 29]. Assembler outputs of each isolate were compared with each other using Mauve to examine circularization of the chromosome and identify any discrepancies, which were resolved with short-read support [30]. To circularize chromosomes, the ends of contigs were manually aligned to determine overlap and were trimmed accordingly. All assemblies were oriented to the same start point to match previously published sequences. To further improve the assemblies, short-read sequences were aligned to the long-read assembly of the same isolate using the vSNP (Step 1); and Pilon with default settings was used to generate consensus base calls [31, 32]. Iterations of alignment and consensus calling were repeated until no SNP changes were observed between iterations. Final assemblies were uploaded to NCBI along with the raw data and annotation with PGAP was completed [33]. Sequencing statistics for all samples (long and short-reads) are in Supplemental Table 2.

### 2.4 Core Genome Determination and Lineage Analysis

Fasta files of de novo, short-read assemblies and complete assemblies were annotated with PROKKA (regardless if NCBI annotations existed) to ensure consistency. Panaroo was used to determine core and pangenomes between all isolates in the study, and a core genome alignment with MAFFT was output. IQTree was used with model selection to generate a phylogenetic tree based on the core genome alignment. This tree was then used to perform lineage analysis with Fast BAPS [34].

### 2.5 NCBI Data

All data used in this study is listed in Supplementary Table 2 and 3 and has been made public through NCBI repositories. For the purposes of developing a more complete phylogeny and predicting GEIs, a dataset defined here as the ‘global dataset’ was used. This dataset was comprised of the NCBI assemblies (7 TE genomes closed for this study and submitted to NCBI for annotation as well as 44 additional *T. equigen-italis* isolates and *T. asinigenitalis* isolate from the NCBI Assembly database) and NCBI Sequence Read Archive short-read data (215 Illumina sequenced isolates and an additional 97 TE and 48 TA isolates sequenced for this study) [33, 35]. To examine the behavior of the GEIs under normal environmental conditions, 141 isolates from a recent US outbreak were used, a dataset defined here as the ‘outbreak dataset’.

### 2.6 Predicting Genomic Islands

IslandCompare was used to search the assemblies in the global dataset for GEIs using the GenBank annotation files from NCBI [36]. IslandViewer4 was then used to generate genome plots of the assemblies closed for this study with GEI locations, and through this process an additional potential GEI was identified (Supplemental Figure 1). Classification and further refinement of the GEIs was performed with ICEberg 3.0s ICEfinder, a tool specifically aimed at classifying integrative and conjugative elements (ICE), integrative and mobilizable elements (IME), and cis-mobilizable elements (CIME) [37]. Each TE GEI in each isolate was compared with others of the same type identified in other isolates. Margins of the GEIs were evaluated to ensure consistency across isolates and were manually adjusted as warranted by homology to genes included in the genomic island in other isolates. GEIs were then extracted and aligned using MAFFT in Geneious Prime version 2022.1.1 to generate identical sites per genomic island [38, 39]. Final versions of each assembled GEI were aligned in pyGenomeViz for visualization (https://github.com/moshi4/pyGenomeViz).

A representative sequence of each GEI was compiled into a single fasta-formatted file for an alignment reference with the short-read sequences in the global dataset. Each short-read sequenced isolate was aligned to the compiled reference fasta with vSNP Step1 and alignment statistics were derived using Samtools [31, 40, 41]. Compiled statistics for the GEIs were evaluated for percent coverage and depth with at least 80% identity, for four of the six *T. equigenitalis* GEI’s (Groups 1-3, and 5), isolates primarily clustered into 2 distinct groups by percent coverage alone. For the 4th and 6th GEIs, there was no clear division in coverage to establish presence or absence. Visual inspection of alignments between isolates predicted to carry a GEI and those without the prediction show variation in a region primarily containing rearrangement hot spot genes. When isolates were evaluated for coverage specifically in this region, two coverage groups were formed. To ensure that the coverage groups correlated with presence/absence of each GEI, short-read sequences of the assemblies (where available) were aligned to the compiled GEI reference. These were then evaluated to establish appropriate coverage cutoffs for inclusion/exclusion. The presence/absence determination based on this short-read data was compared with predictions on the assemblies to ensure consistency in predictions.

### 2.7 Stability of Genomic Islands

The outbreak dataset was used to examine the stability of GEIs within the genome in the natural environment. Caution was used to avoid serial passaging of cultures prior to DNA extraction, which attempted to avoid laboratory adaptation and the possible loss of GEIs under laboratory conditions. Sequences were aligned to the GEI reference in the fasta format used previously, and isolates were characterized for the presence and absence of each genomic island as described above. To illustrate the genetic relatedness of isolates, vSNP was used with reference NZ CP0076337 to generate a SNP-based phylogeny [31]. Initially, the tree was built using all SNP data including GEIs; and a second tree was generated with SNP-data within the GEIs removed to visualize genetic relatedness without the influence of the mobile elements.

## 3 Results

### 3.1 Phylogenetics

The kSNP tree of both *Taylorella* species shows distinct separation between *T. equigenitalis* and *T. asinigenitalis* (Figure 1A); and, within the *T. equigenitalis* tree, the isolates are sorted into highly divergent clades (Figure 1B). Overall, the trees show significant genetic diversity while clustering outbreak isolates as well as some isolates with common geographic origins. The long, internal branches are consistent with results of the recent Czech study which places the most recent common ancestor of *T. equigenitalis* around 640 A.D. (80-900 A.D.) [9]. The 7 closed genomes from this study provide the first closed reference for 6 clades as well as re-sequencing of MCE9.

**Fig. 1.**
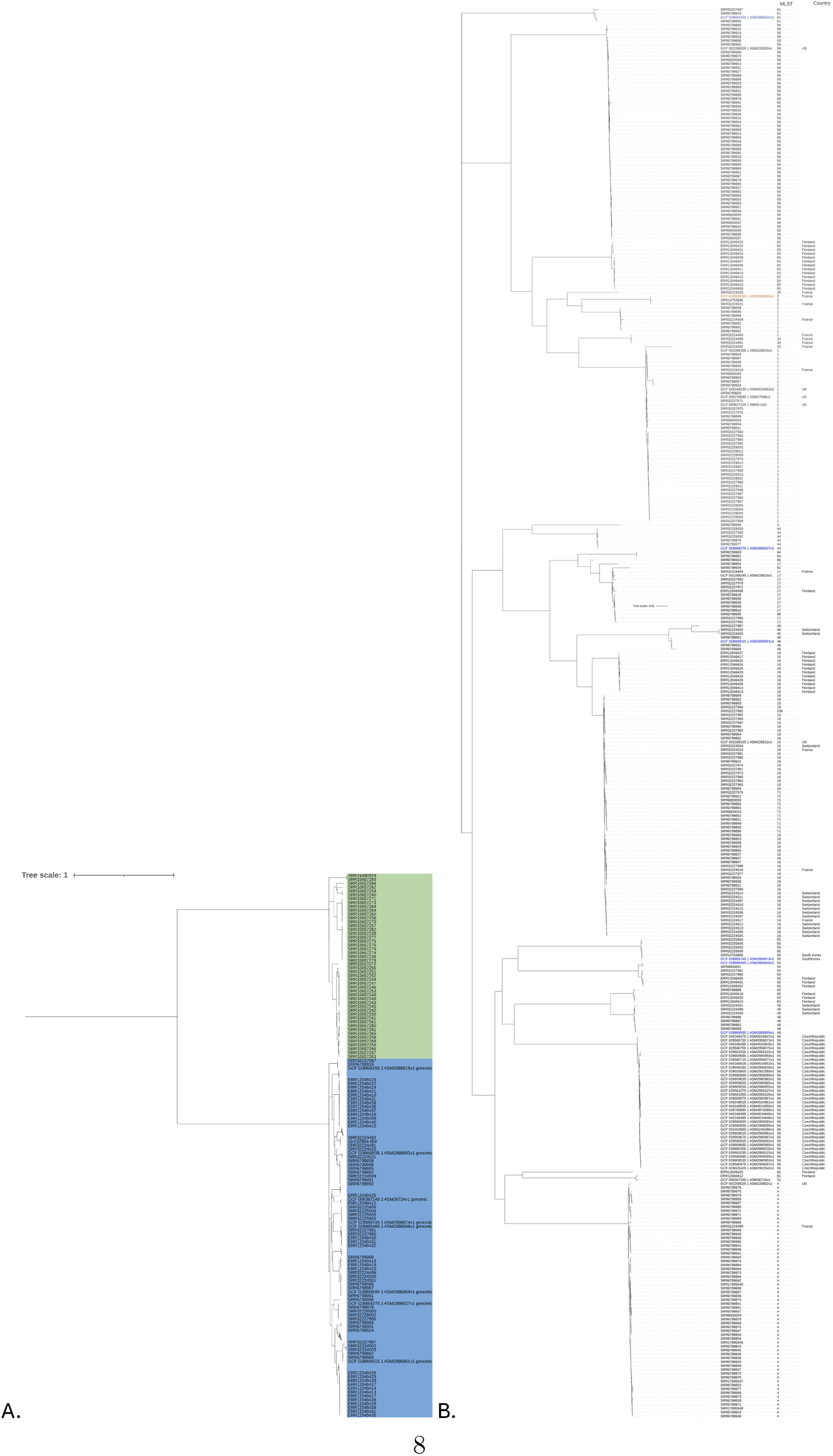
A. Phylogenetic tree of *T. asinigenitalis* (green) and *T. equigenitalis* (blue). *T. equigenitalis* clades with large numbers of isolates have been collapsed for viewing purposes. B. Phylogenetic tree of *T. equignitalis* with MLSTs and country of origin noted. Isolates without countries listed are isolates from animal movement cases where a single country could not be attributed. Isolates in blue represent genomes closed for this project, and in orange is MCE9, which was re-sequenced and closed for this project.

### 3.2 Core Genome and Lineage Analysis

The core genome (*≥* 99% of genomes) contained 1404 genes, 70.5% of the total genes identified in *T. equigenitalis*. Alignment of these genes and subsequent lineage analysis yielded 7 main lineages that further divided into 15 sub-lineages. These occurred along major branches of the core genome phylogenetic tree (Figure 2).

**Fig. 2.**
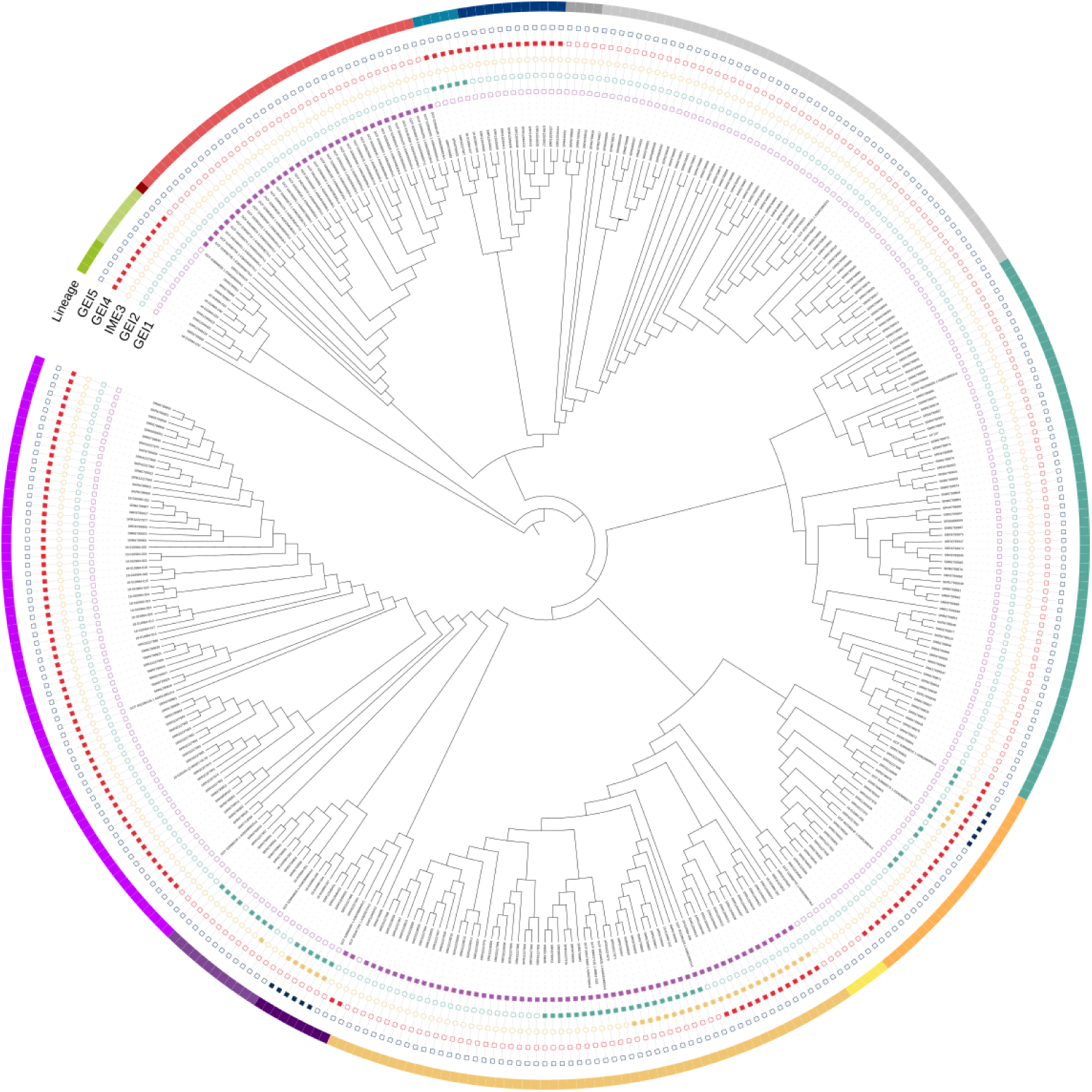
Core genome phylogenetic tree. This tree, generated by IQTree, shows the 2-level lineage analysis and presence of GEIs 1-5. Each level 1 lineage is denoted by color category with shading representing level 2 sublineages.

### 3.3 Genomic Islands

A total of 7 GEIs were identified with 6 being found in only *T. equigenitalis* genomes and 1 in the *T. asinigenitalis* genome. IslandCompare identified 5 unique GEIs (named GEI1 to GEI5) across all 51 *T. equigenitalis* assemblies with each genome having 0-4 GEIs (Supplemental Table 3). IslandViewer4, based on 12 assemblies completed by the NVSL, predicted the same GEIs as Island Compare plus a potential 6th *T. equigenitalis* GEI (named GEI6) (Supplemental Figure 1). ICEberg’s ICEfinder refined the positions of 2 GEIs, GEI2 and GEI3, and provided classifications of both. Regardless of prediction tool, GEIs of the same group show a high degree of similarity across genomes as seen in Table 1. Annotations of each GEI are shown in Supplemental Table 1.

**Table 1.**
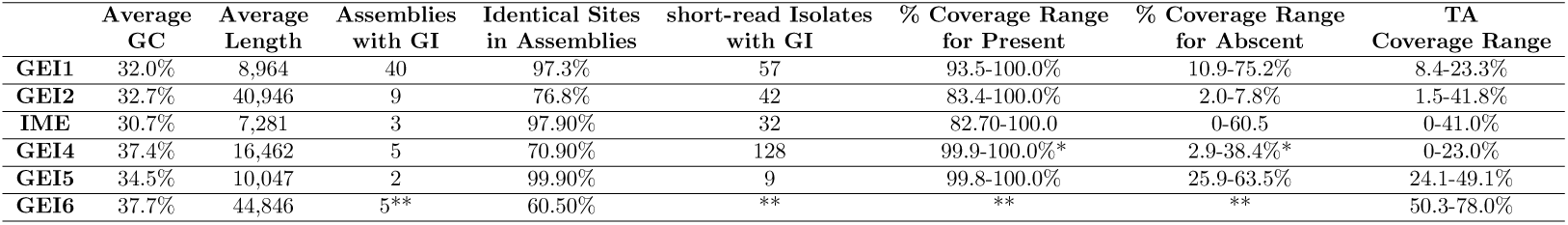
Number of Isolates with GEI.

Evaluated from position 1000 to 2700

Only the 12 assemblies from the NVSL were included in this prediction. Presence/Absence metrics could not be determined.

#### 3.3.1 Genomic Island 1

GEI1 was the most prevalent GEI in the assemblies, occurring in 40 of the 51 *T. equigenitalis* assemblies evaluated. However, this may be due to sampling bias as 35 of the 40 assemblies containing this GEI occur in a single clade. All 40 assemblies represent only 4 clades labeled in the *T. equigenitalis* tree (Figure 3). GEI1 remained highly conserved across the 40 isolates with only 6 unique sequences observed and 98.1% identical sites between them. The 6 sequences can only be differentiated by SNPs and small insertions and deletions, but annotations (Supplemental Table 1) remained the same between all the sequences. The initiation of this GEI was always located between 1.28 Mb and 1.35 Mb with placement being obviously influenced by the presence of other GEIs.

**Fig. 3.**
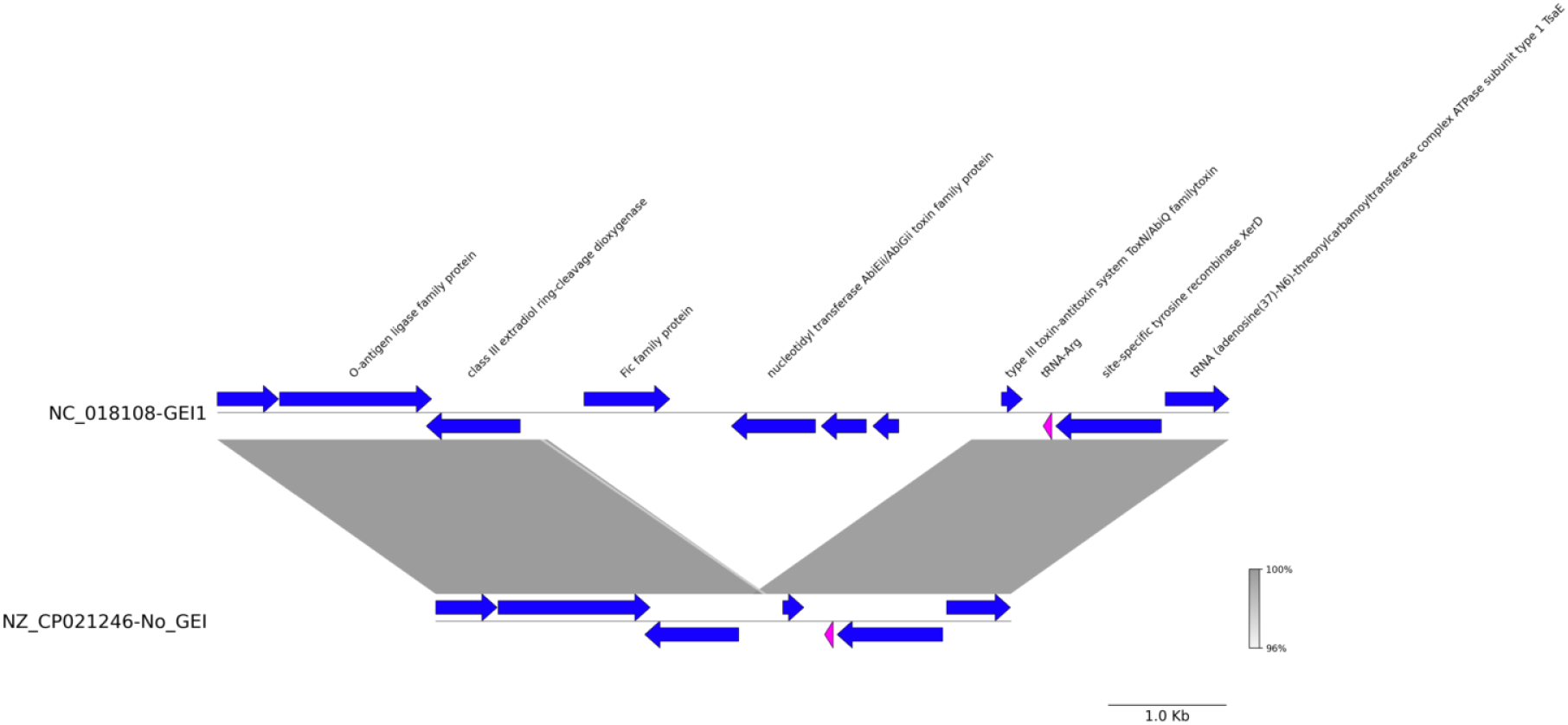
Alignment of GEI1. GEI1 in isolate NC_018108, while isolate NZ_CP021246 is representative of TE isolates without the GEI predicted. Genes without annotation are hypothetical. Aligned segments represent 96% or greater BLAST matches.

Among the *T. equigenitalis* short-reads, coverage statistics for GEI1 primarily fell into 2 groups: 97-100% and 53-75% and only 1 isolate had less than 53% coverage. Fifty-seven isolates carrying the GEI1, were in the 97-100% group with an overall average depth of coverage of 94X, making this the second most common GEI identified in the short-reads. However, the predicted GEI was rarely entirely absent in any isolates examined. As shown in Figure 3, a truncated GEI1-like sequence was observed in 261 isolates, which comprised the group with 53-75% coverage. These isolates had an average depth of coverage of 82X and contained a deletion within GEI1 which spanned from a gene enconding the Fic family protein to the hypothetical protein located at the 3’ end of the toxN gene.

Conversely, alignment of the *T. asinigenitlais* short-reads to this same reference showed low coverage at 8-23% but greater than 90% identity in the covered regions.

#### 3.3.2 Genomic Island 2

This is the most widely dispersed GEI in the *T. equigenitalis* tree (Figure 2), being found in isolates from 7 of the 13 major clades. GEI2 is the largest of all the GEIs identified in this study, classified as a T4SS-ICE by ICEberg. The 9 assemblies containing this GEI showed variability, with 5 different annotation patterns (Figure 4). Notably, 3 isolates lacked a contiguous genomic region near the 3’ end containing the ToxN/AbiQ family toxin gene, the repA gene, and a helix-turn-helix transcriptional regulator. Despite this 2,556 bp gap, the GEI is composed of 76.8% identical sites across the 9 isolates, and it was consistently located within genomes with the 5’ end of GEI2 always located between 1.17 Mb and 1.21 Mb in the genome. Among the short-reads, the coverage statistics formed 2 distinct groups: 83%-100% and less than 10%, with 47 isolates falling into the first group and 272 isolates in the second group.

**Fig. 4.**
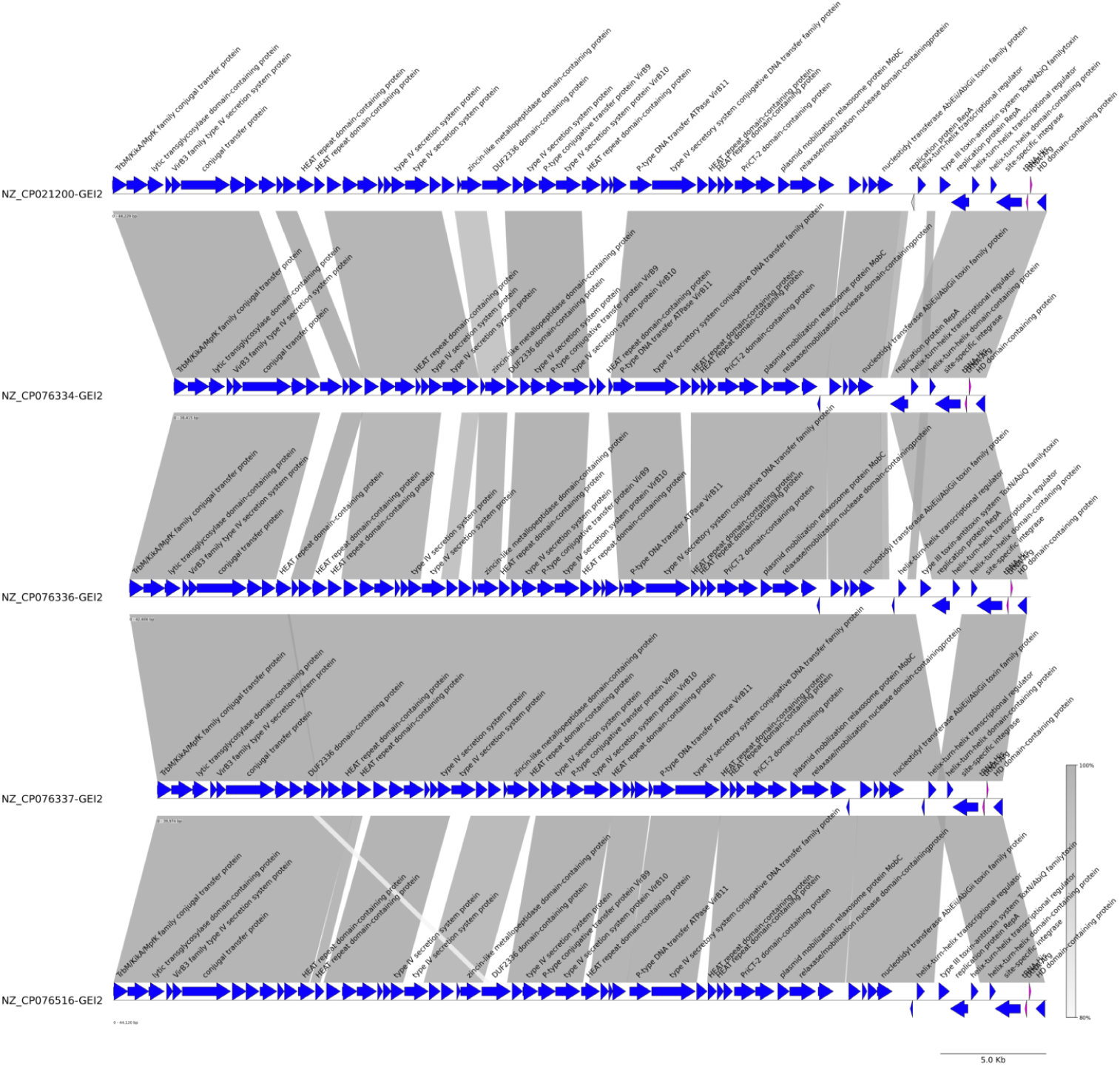
Alignment of GEI2. Among the assemblies with this genomic island, 5 variations in the annotations were found. The alignment of these sequences and annotations shows overall homology with discrete changes in the island. Genes without annotation are hypothetical. Aligned segments represent 80% or greater BLAST matches.

Among the *T. asinigenitalis* isolates, average coverage ranged from 2 to 42% in isolates with 37 of the 47 isolates having 5.3% or less coverage. All *T. asinigenitalis* iso-lates had greater than 90% identity within the coverage regions, regardless of percent coverage.

#### 3.3.3 Genomic Island 3

GEI3 was predicted in 4 annotated genomes by IslandCompare. When these genomes were analyzed in ICEberg’s ICEfinder, a shifted, more refined region that included the site-specific integrase and AAA family ATPase immediately upstream of the IslandCompare prediction was classified as an integrated mobilizable element (IME). However, the integrase is not present in 1 of the assemblies, leaving only 3 assemblies with the IME identified (Figure 5). As with the previous GEIs, it was consistently located in the same region of each genome beginning between 0.97-0.98 Mb in the genome. Of the short-reads, 34 had 98% or greater coverage to the IME. One additional short-read isolate did not hae the site-specific integrase (83% coverage). There was a significant drop in coverage in the remainder of the isolates - primarily *≤* 10%, with only 2 isolates showing slightly higher values at 14% and 17%.

**Fig. 5.**
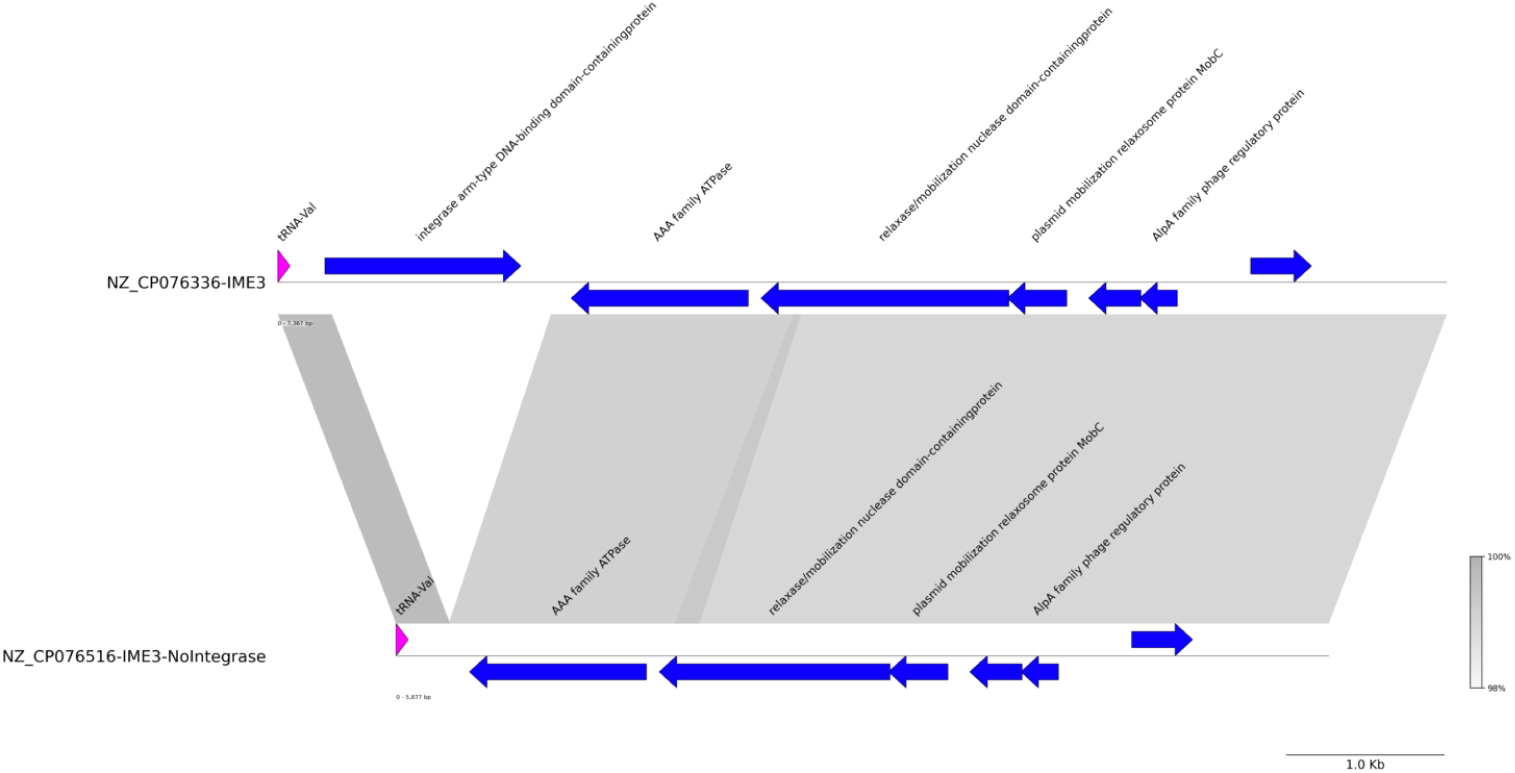
Alignment of Sequences Representing IME3, and No Genomic Island Predicted. NZ_CP076336 contained the full IME3, while NZ_CP076516 is missing the integrase within the IME. Genes without annotation are hypothetical. Aligned segments represent 98% or greater BLAST matches.

Most *T. asinigenitalis* isolates had 10% or less coverage of this GEI, but 4 isolates had 21%-25% coverage, and a single isolate had 41% coverage. While percent identity varied widely among isolates with the lowest coverage, the 5 isolates with higher coverage all had 89% or greater identity in coverage regions.

#### 3.3.4 Genomic Island 4

GEI4 had greater than 85% coverage in nearly all assemblies and short-read sequences. The predominant region of variability occurred near the 5’ end in a set of genes annotated as hypothetical protein and Rearrangement Hot Spots, which appears to have separated assemblies with the predicted GEI from those without. Outside of this region, isolates contained genes encoding a transposase as well as TssI/VgR, ATP binding protein, CstA (pyruvate/proton symporter), TcuAB (tricarballylate utilization 4Fe-4S protein and tricarballylate dehydrogenase). The *tcuA* gene contained a PAAR domain (proline-alanine-alanine-arginine), an essential counterpart to Tssi/VgrG to form the VgrG spike of the type VI secretion system (T6SS) [13].

In the assemblies, 5 were predicted to contain this GEI, each having some variation in the annotation of the region as seen in Figure 6. Among the short-reads, when evaluated in the limited region of diversity near the 5’ end (as described in the methods), 131 isolates were predicted to carry GEI4. This GEI consistently appeared in the genomes between positions 1.45 Mb and 1.51 Mb. It was also widely dispersed in the phylogenetic tree, occurring in nearly every cluster of isolates throughout as seen in Figure 3.

**Fig. 6.**
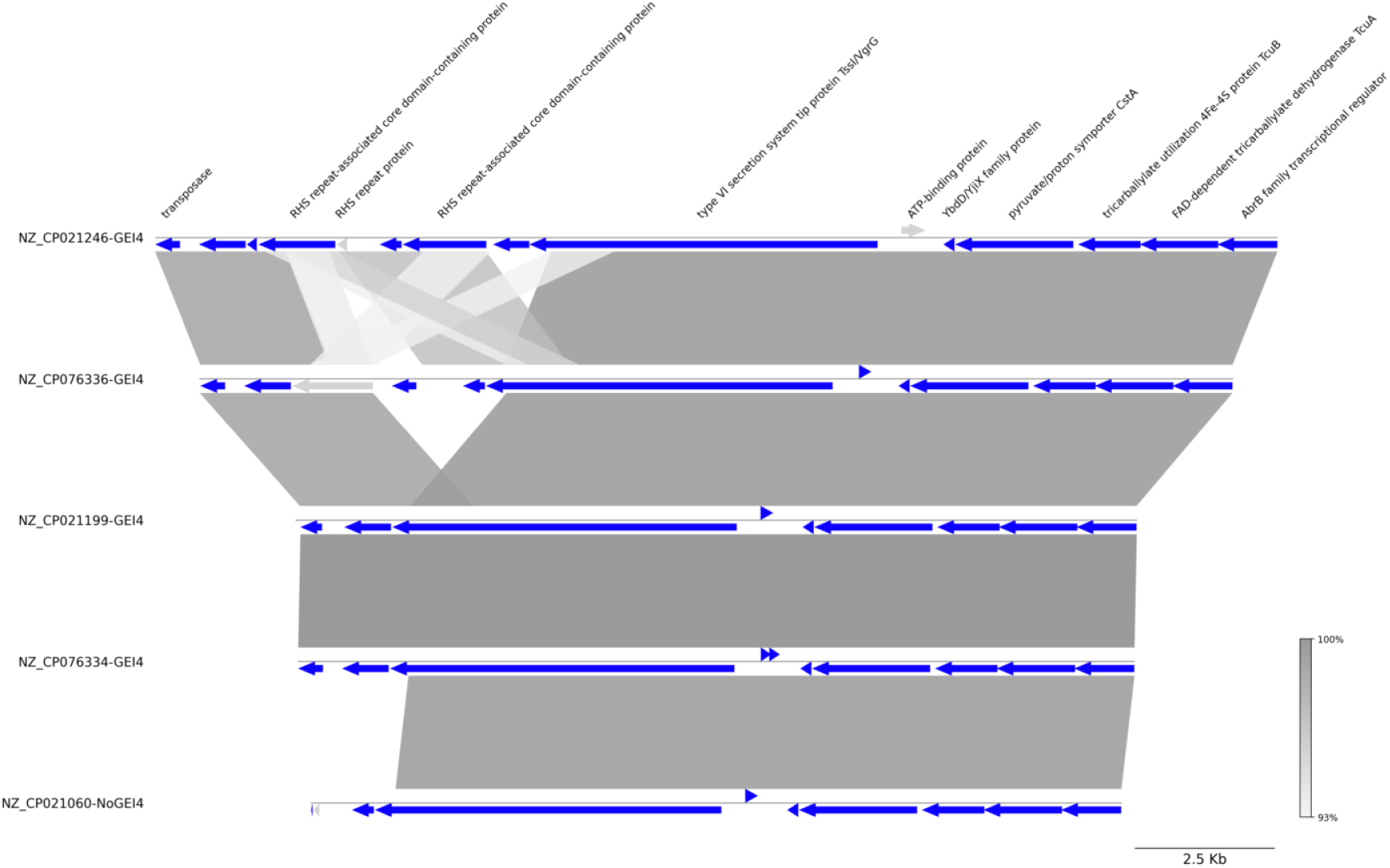
Alignment of GEI4 Sequences with a Sequence Found in the Same Region Not Predicted to be GEI4. Four annotation variations were observed for this GEI. NZ_CP021060 contains a sequence in the same region of the genome and with homology to a portion of GEI4 observed in other genomes. Genes without annotation are hypothetical. Aligned segments represent 93% or greater BLAST matches.

Among the *T. asinigenitalis* isolates there was 32% - 53% coverage across the entire GEI and all isolates had 92% or great percent identity.

#### 3.3.5 Genomic Island 5

GEI5 was predicted in only 2 assemblies by IslandCompare and was also the least common GEI in the short-reads, with only 11 isolates having more than 99% coverage while the remainder of the short-read sequences had less than 61% coverage. Alignment of the annotated region between the two coverage groups is visible in Figure 7. Like the other GEIs, this was also consistently located in the genomes beginning near 0.91 Mb. The dispersion in the *T. equigenitalis* tree can be seen in Figure 2 and is expectedly limited.

**Fig. 7.**
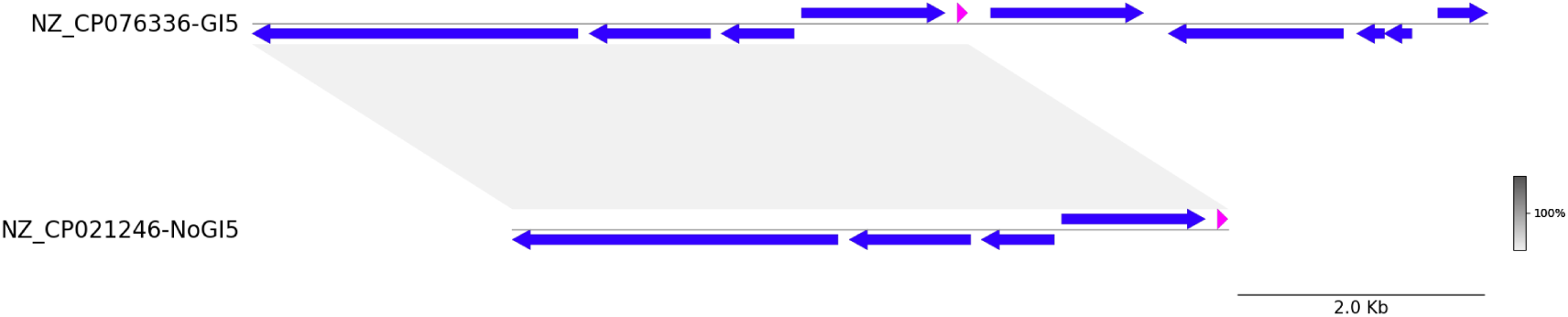
Alignment of GEI5 with Sequence Commonly Located in the Same Region. The bottom sequence was predicted to be a genomic island by IslandCompare and IslandViewer4, while the bottom sequence was not and is representative of most TE sequences. Genes without annotation are hypothetical. Aligned segments represent 80% or greater BLAST matches.

*T. asinigenitlais* isolates had 24% - 49% coverage of this region with 88% or greater percent identity.

#### 3.3.6 Potential Genomic Island 6

IslandViewer4 identified a potential sixth GEI in 5 of the 12 assemblies. Among the 5 assemblies, a MAFFT alignment indicated 61% identity. Despite the low percent identity, the annotations between these 5 isolates in the identified region were highly similar. However, they were also highly similar to isolates where the GEI was not predicted. Extraction of homologous regions from all the assemblies revealed similar annotations across all isolates with nucleotide variation limited to discrete segments. No specific variation or pattern was linked solely to isolates predicted to have GEI6, preventing differentiation between predicted presence and absence. Additionally, all short-read *T. equigenitalis* isolates had 86% or more coverage in this region and 96% identity or more, while *T. asinigenitalis* isolates had 50-78% coverage of this same region with 93% or greater identity of the covered regions. This represents the only prediction in the *T. equigenitalis* isolates to also have significant coverage in the *T. asinigenitalis* genomes. Annotations of the region are outlined in Supplemental Table 1, with observed variations depicted in Figure 8. The list of annotations from all variations is available in Supplemental Table 4. These annotations included TssI/VgrG proteins, immunity proteins, RHS domains, ankyrin repeat domain-containing protein, and various proteins with enzyme domains. Further analysis identified that the NAD-dependent succinate-semialdehyde dehydrogenase is a PAAR domain-containing protein.

**Fig. 8.**
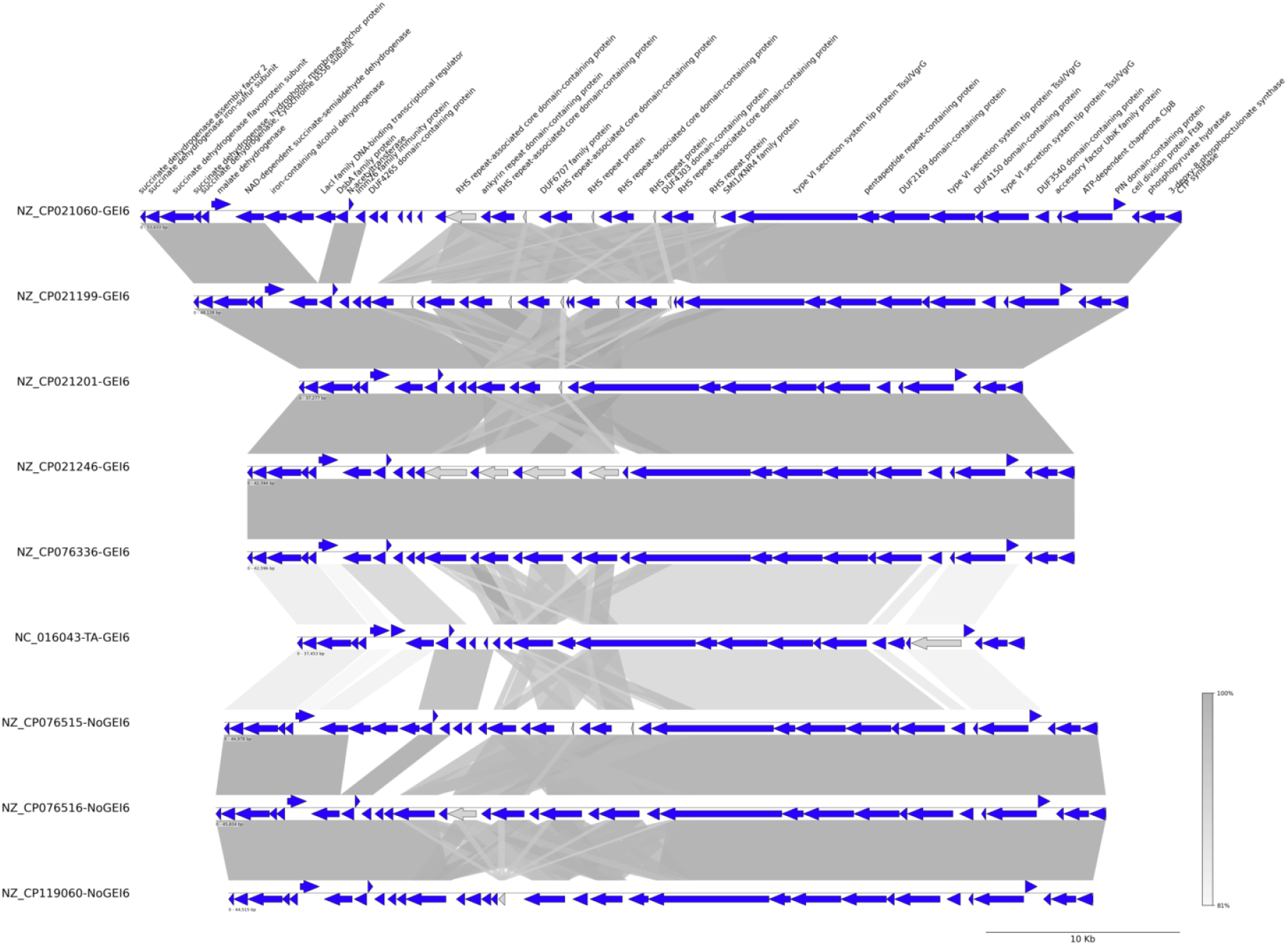
Alignment of Potential GEI6. Included are all 5 genomes where GEI6 was predicted, the homologous region from TA isolate MCE3, and homologous regions from several assemblies where the GEI was not predicted. Genes without annotation are hypothetical. Aligned segments represent 81% or greater BLAST matches.

Annotations are listed in the order they appear, but when multiple hypothetical protein CDS were observed in a row, they were abbreviated with the number of times observed in order to abbreviate the table.

#### 3.3.7 *T. asinigenitalis* Genomic Island

The GEI in the *T. asinigenitalis* isolate was distinct from the *T. equigenitalis* GEIs. It was the largest GEI identified in this study, with 69 annotated genes across 53,195 bp (Table 2). Of these, 41 encoded hypothetical proteins, making it difficult to elucidate the function of this GEI. Of the genes that were annotated, 3 genes appear in duplicate including ninB, ydaS, and a gene encoding an S24 family peptidase.

**Table 2.** Gene Annotations.

| Gene/Protein |
| --- |
| GNAT family N-acetyltransferase CDS |
| NAD(P)H-quinone oxidoreductase CDS |
| tpiA CDS |
| secG CDS |
| tRNA-Leu |
| site-specific integrase CDS |
| **2 x** hypothetical protein CDS |
| DNA adenine methylase CDS |
| helix-turn-helix domain-containing protein CDS |
| ssb CDS |
| hypothetical protein CDS |
| phage protein CDS |
| hypothetical protein CDS |
| siphovirus Gp157 family protein CDS |
| ATP-binding protein CDS |
| **3 x** hypothetical protein CDS |
| S24 family peptidase CDS |
| YdaS family helix-turn-helix protein CDS |
| recombination protein NinB CDS |
| hypothetical protein CDS |
| ParB N-terminal domain-containing protein CDS |
| DEAD/DEAH box helicase family protein CDS |
| toprim domain-containing protein CDS |
| DUF1064 domain-containing protein CDS |
| **7 x** hypothetical protein CDS |
| terminase small subunit CDS |
| terminase family protein CDS |
| **12 x** hypothetical protein CDS |
| DUF6682 family protein CDS |
| **7 x** hypothetical protein CDS |
| glycoside hydrolase family 104 protein CDS |
| LPD38 domain-containing protein CDS |
| hypothetical protein CDS |
| lysozyme CDS |
| hypothetical protein CDS |
| DUF1902 domain-containing protein CDS |
| S24 family peptidase CDS |
| YdaS family helix-turn-helix protein CDS |
| recombination protein NinB CDS |
| **5 x** hypothetical protein CDS |

The coverage of this GEI varied in the TA short-read isolates, 2 samples had 92% coverage, 3 had approximately 41% coverage, and the remainder of the isolates had *≤* 26%coverage of the GEI. *T. equigenitalis* coverage was low for all samples in this study, with a maximum of 16% and 222 isolates having *≤* 2% coverage.

### 3.4 Stability of Genomic Islands

To illustrate the dispersion of GEIs in outbreak isolates and their influence of phylogenetic structure, SNP based phylogenetic trees are shown in Figure 9. A phylogenetic tree with the SNPs in the GEIs included can be seen in Figure 9A, while Figure 9B has SNPs in the GEIs filtered to remove their influence on the tree structure. Using the cutoffs established above, 100% of this outbreak dataset contained partial coverage of GEI1 and potential GEI6, with GEI1 ranging from 57-76% coverage, which was consistent with the results seen from isolates in the global dataset described above. Potential GEI6 ranged in coverage from 89-90%, also consistent with the results seen in isolates from the global databaset. GEI2, IME3, and GEI5 were detected in 42%, 85%, and 36% of the outbreak dataset, respectively, while no isolates contained GEI4. It is of interest to note that 4% of these isolates contained no GEIs and 22% contained all 3 GEIs identified. The remainder of isolates were distributed between different patterns of GEIs with the exception of only GEI5, which wasn’t observed in any isolates. Table 3 shows the patterns of GEIs observed.

**Fig. 9.**
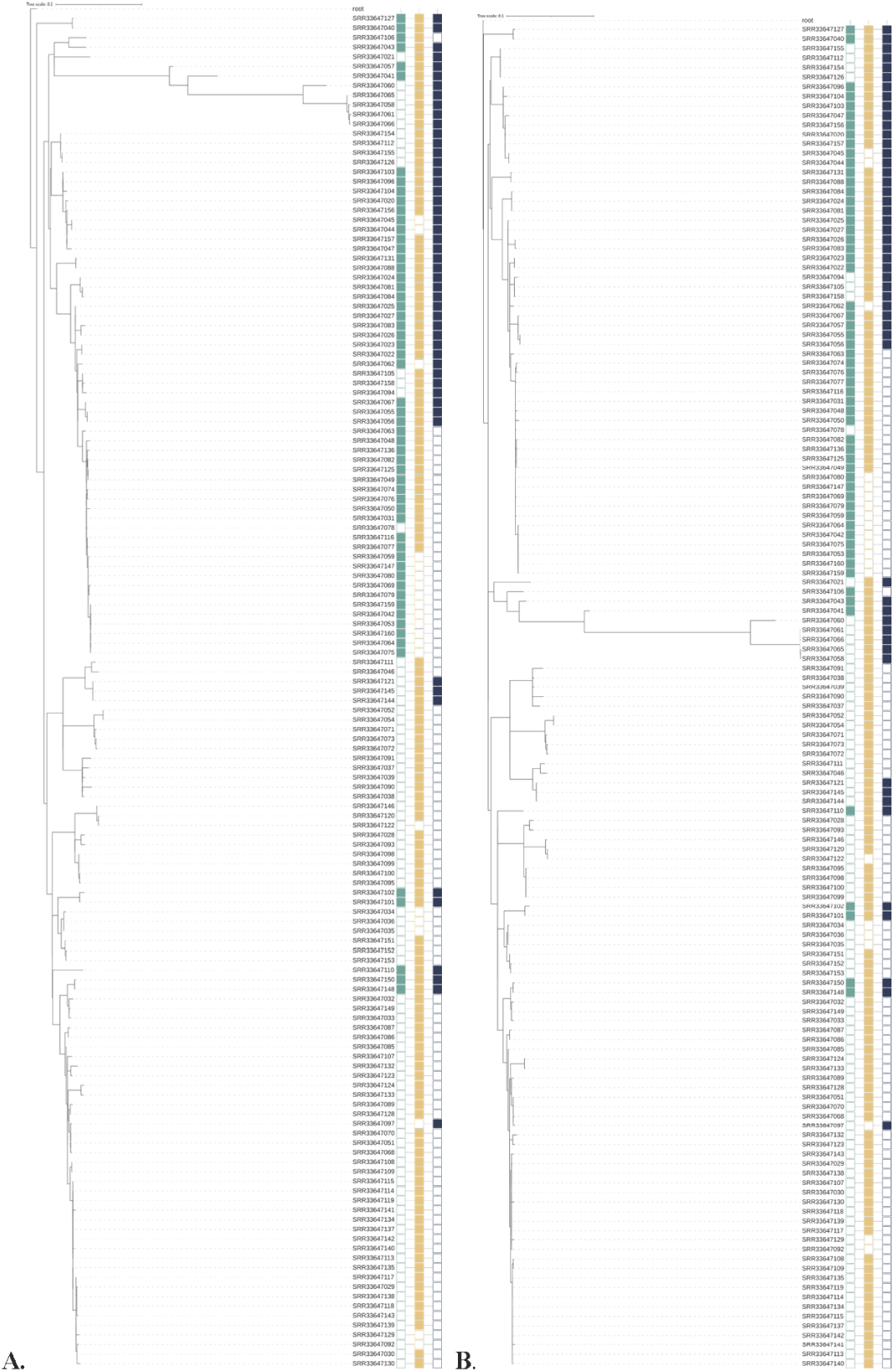
A) is based on all high-quality SNPs in all of the genomes. B) is based on SNPs outside the GEIs only.

**Table 3.** Occurrence of GEIs in the Outbreak Isolates.

| Pattern | Partial GEI1, GEI2, IME3, GI5 | Partial GEI1, GEI2, IME3 | Partial GEI4, GEI2, GEI5 | Partial GEI1, IME3, GEI5 | Partial GEI1 & GEI2 Only | Partial GEI1 & IME3 Only | Partial GEI1 & GEI5 Only | Partial GEI1 ONLY |
| --- | --- | --- | --- | --- | --- | --- | --- | --- |
| Count | 31 | 13 | 3 | 17 | 12 | 60 | 0 | 6 |
| Percentage | 22% | 9% | 2% | 12% | 8% | 42% | 0 | 4% |

## 4 Discussion

*T. equigenitalis* is of concern in equine trade due to its potential for causing disease, often resulting in acute infertility in mares and economic loss in the breeding industry. However, the severity of clinical signs can vary widely, ranging from undetectable to severe. To better predict strain virulence, it is essential to characterize virulence factors and potential recombination mechanisms. By combining closed genomes with short-read sequencing data from isolates in this study, we constructed a more accurate phylogenetic framework for *T. equigenitalis*. This comprehensive genomic analysis enables systematic comparisons of clade-specific characteristics and provides critical foundational data for developing improved molecular diagnostics and epidemiological tracking systems for this pathogen.

The GEIs identified in this study mark the first identification of mobile genetic elements in Taylorella spp. and represent the potential for recombination between genomes. As in other organisms, these GEIs demonstrate that conserved sequences across many branches of the phylogenetic tree, occur at discrete locations within the genomes, and possess the genetic mechanisms for mobility; however, this genus contains many hypothetical proteins. Functional understanding of these genes may elucidate additional roles of these GEIs. The genes that are annotated (or contain a functional domains) represent potential functions in virulence and ecological fitness.

GEI2 contains virB3 and virB8-11, often associated with transcription of virulence factors as well as being components of T4SSs [13, 42]. Additional T4SS proteins without further classification were also present. Among the annotated genes and functional domains beyond the T4SS were an AbiEii/AbiGii toxin family protein as well as a type III toxin-antitoxin ToxN/AbiQ family protein, also seen in GEI1. Noteably, plasmid mobilization protein mobC was also present along with 2 conjugal transfer proteins, site-specific integrase, RepA, and ArdC and AlpA family proteins.

Many of these proteins are known to be involved in mobilization and conjugation, while others are anti-defense mechanisms [43, 44]. The AlpA phage regulatory protein has been associated with regulation of cell lysis [45]. T4SSs have been associated with virulence and organism ecology, and the genes found here appear to support these associations [22].

IME3 and GEI4 are less unique. Like GEI2, IME3 contains mobC and AlpA (also found GEI5), making its contents duplicative of other GEIs. The GEI4 region primarily appears to consist of genes involved in metabolism and possibly catabolism with the clear exemption of tssI/vgrG, a T6SS spike protein and the associated PAAR domain (like GEI6) [46–48].

GEI5 annotations in Supplemental Table 1 include genes spanning a range of functions. In addition to the AlpA disussed previously, also present is a TRAP transporter, annotated as a fused permease subunit and a solute-binding subunit, is commonly associated with nutrient uptake from the environment; while the AI-2E transporter belongs to a superfamily that has little published data, but has been associated with signaling pathways [49, 50].

GEI4 and the potential GEI6 share structural similarities, particularly in their use of RHS domains and the absence of tRNAs. They also do not deviate from the GC content of the whole genome unlike other GEIs [51]. Notably, the T6SS protein domain TssI, the spike protein, is found in both these GEIs with accompanying PAAR domains. These domains are highly conserved across all isolates, regardless of GEI predictions, and are often virulence determinants in organisms[13]. In the potential GEI6, several immunity protein domains are also conserved, extending even across the genus, which may preclude this region from being a genomic island, as they are generally species specific. However, these immunity proteins are associated with polymorphic immunity systems or heterogenous polyimmunity loci. In these systems the spike protein is a common method for delivering bactericidal effectors into neighboring cells with the help of immunity proteins that prevent the cell’s own death. These systems clearly enhance an organism’s virulence with the ability to deliver toxins and/or effectors into neighboring cells [52–54].

Many of the genes within these predicted GEIs are likely to have additional functions, particularly anti-defense, aiding this fastidious organism in surviving and thriving within competitive environments, such as the male reproductive tract. However, the data set includes only a limited number of isolates that have been characterized in terms of clinical signs, virulence levels, and other phenotypic traits. This limitation prevents any definitive correlation between the GEIs identified in this study and phenotypic data. Further research is essential to determine whether these GEIs are linked to environmental stressors or variations, and to explore how they might impact virulence or enhance the organism’s survivability.

The recent U.S. outbreak of CEM, which resulted from a single introduction with clonal expansion, according to molecular epidemiological studies as well as epidemiological tracing, provided an opportunity to examine GEIs identified in this study for potential mobility under natural conditions. This outbreak has occurred primarily among geldings through fomite transmission with numerous opportunities for re-exposure of animals over a period of several years, limiting the conclusions that can be drawn from this data beyond demonstrating the organism’s ability to excise or acquire these GEIs. The various patterns observed may well be correlated with usefulness of the genes contained in the GEIs present and their ability to improve survivability of the *T. equigenitalis* strain.

Further investigation into the functional potential of these genomic islands could enlighten our understanding of *T. equigenitalis*’s pathogenicity and potentially reveal insights into the pathogenic potential of *T. asinigenitalis*.

## 5 Conclusion

This work demonstrates several genomic islands, their associated genes, and their distribution throughout a more complete phylogenetic tree. This is the first documentation of mobile genetic elements in this genus.

## Supplementary information

- Supplemental Figure 1. IslandViewer4 Output graphs of the 12 *T. equigenitalis* assemblies from the NVSL.
- Supplemental Table 1. *T. equigenitalis* Genomic Island Annotations
- Supplemental Table 2. Assembly stats and raw read accessions for samples assembled in this study.
- Supplemental Table 3. NCBI SRA Accession Numbers
- Supplemental Table 4. Presence/Absence of GEIs in Assemblies
- Supplemental Table 5. Annotation list from Potential GEI6 of each assembly

## Acknowledgements

We would like to thank Ian Mawhinney and Animal and Plant Health Agency, Veterinary Investigation Laboratory, Suffolk, United Kingdom for contributing isolates that expanded the phylogenetic trees with new clades. His insightful questions and discussions on isolate variation helped shape the concept of this study. We also thank Dr. Alan Guthrie, Equine Research Centre, University of Pretoria, South Africa for providing isolates and sharing his expertise on T. equigenitalis. This work would not have been possible without their valuable contributions.

## 6 Declarations

- No funding was received for this work.
- The author(s) declare(s) that they have no competing interests.
- All data used in this study is available through NCBI. A full list of accession numbers for assemblies and raw data is available in Supplemental tables 2 and 3.
- Author contribution

## References

[1] Platt, H., Atherton, J.G., Simpson, D.J., Taylor, C.E., Rosenthal, R.O., Brown, D.F., Wreghitt, T.G.: Genital infection in mares. Veterinary Record 101, 20 (1977)

[2] Taylor, C.E.D., Rosenthal, R.O., Brown, D.F.J.: The causative organism of contagious equine metritis 1977: Proposal for a new species to be known as haemophilus equiqenitalis. Equine Vet. J. (1978)

[3] Platt, H., Atherton, J.G.: The experimental infection of ponies with contagious equine metritis. Equine Vet. J. (1978)

[4] Swerczek, T.W.: The first occurence of contagious equine metritis in the united states. JAVMA 173(4), 405–407 (1978)

[5] Erdman, M.M., Creekmore, L.H., Fox, P.E., Pelzel, A.M., Porter-Spalding, B.A., Aalsburg, A.M., Cox, L.K., Morningstar-Shaw, B.R., Crom, R.L.: Diagnostic and epidemiologic analysis of the 2008-2010 investigation of a multi-year outbreak of contagious equine metritis in the united states. Prev Vet Med 101(3-4), 219–28 (2011) 10.1016/j.prevetmed.2011.05.015

[6] Bleumink-Pluym, N.M.C., Lakk, E.A., Van Der Zeust, B.A.M., Houwers, D.J.: Differences between taylorella equigenitalis strains in their invasion of and replication in cultured cells. Clinical and Diagnostic Laboratory Immunology (1996)

[7] Aalsburg, A.M., Erdman, M.M.: Pulsed-field gel electrophoresis genotyping of taylorella equigenitalis isolates collected in the united states from 1978 to 2010. J Clin Microbiol 49(3), 829–33 (2011) 10.1128/JCM.00956-10

[8] Duquesne, F., Hebert, L., Breuil, M.F., Matsuda, M., Laugier, C., Petry, S.: Development of a single multi-locus sequence typing scheme for taylorella equigenitalis and taylorella asinigenitalis. Vet Microbiol 167(3-4), 609–18 (2013) 10.1016/j.vetmic.2013.09.016

[9] Hrala, M., Andrla, P., Bosak, J., Fedrova, P., Mugutdinov, A., Karpiskova, R., Nedbalcova, K., Raichova, J., Faldyna, M., Horin, P., Smajs, D.: Whole genome sequences of nine taylorella equigenitalis strains isolated in the czech republic between 1982-2021: Molecular dating suggests a common ancestor at the time of roman empire. PLoS One 20(1), 0315946 (2025) 10.1371/journal.pone.0315946

[10] Hauser, H., Richter, D.C., Tonder, A., Clark, L., Preston, A.: Comparative genomic analyses of the taylorellae. Vet Microbiol 159(1-2), 195–203 (2012) 10.1016/j.vetmic.2012.03.041

[11] Hebert, L., Moumen, B., Pons, N., Duquesne, F., Breuil, M.F., Goux, D., Batto, J.M., Laugier, C., Renault, P., Petry, S.: Genomic characterization of the taylorella genus. PLoS One 7(1), 29953 (2012) 10.1371/journal.pone.0029953

[12] Hicks, J., Stuber, T., Lantz, K., Erdman, M., Robbe-Austerman, S., Huang, X.: Genomic diversity of taylorella equigenitalis introduced into the united states from 1978 to 2012. PLoS One 13(3), 0194253 (2018)

[13] Wilson, B.A., Winkler, M.E., Ho, B.T.: Bacterial Pathogenesis: A Molecular Approach, 4th edn. (2019)

[14] Green, E.R., Mecsas, J.: Bacterial secretion systems: An overview. Microbiol Spectr 4(1) (2016) 10.1128/microbiolspec.VMBF-0012-2015

[15] Meade, B.J., Timoney, P.J., Donahue, J.M., Branscum, A.J., Ford, R., Rowe, R.: Initial occurrence of taylorella asinigenitalis and its detection in nurse mares, a stallion and donkeys in kentucky. Prev Vet Med 95(3-4), 292–6 (2010) 10.1016/j.prevetmed.2010.04.010

[16] Dorrego, A., Herranz, C., Perez-Sancho, M., Camino, E., Gomez-Arrones, V., Carrasco, J.J., De Gabriel-Perez, J., Serres, C., Cruz-Lopez, F.: First report and molecular characterization of cases of natural taylorella asinigenitalis infection in three donkey breeds in spain. Vet Microbiol 276, 109604 (2023) 10.1016/j.vetmic.2022.109604

[17] Jang, J.S., Donajue, J.M., Arat, A.B., Goris, J., Hansen, L.M., Earley, D.L., al.: Taylorella asinigenitalis sp. nov., a bacterium isolated from the genital tract of male donkeys (equus asinus) (2001) 10.1099/00207713-51-3-971

[18] Wilsher, S., Omar, H., Ismer, A., Allen, T., Wernery, U., Joseph, M., Mawhinney, I., Florea, L., Thurston, L., Duquesne, F., Petry, S.: A new strain of taylorella asinigenitalis shows differing pathogenicity in mares and jenny donkeys. Equine Vet J 53(5), 990–995 (2021) 10.1111/evj.13382

[19] Katz, J.B., Evans, L.E., Hutto, D.L., Schroeder-Tucker, L.C., Carew, A.M., Donahue, J.M., Hirsh, D.C.: Clinical, bacteriologic, serologic, and pathologic fea-tures of infections wtih atypical taylorella equigenitalis in mares. Journal of the American Veterinary Medical Association 12, 1945–1948 (2001)

[20] Dobrindt, U., Hochhut, B., Hentschel, U., Hacker, J.: Genomic islands in pathogenic and environmental microorganisms. Nat Rev Microbiol 2(5), 414–24 (2004) 10.1038/nrmicro884

[21] Bertelli, C., Tilley, K.E., Brinkman, F.S.L.: Microbial genomic island discovery, visualization and analysis. Brief Bioinform 20(5), 1685–1698 (2019) 10.1093/bib/bby042

[22] Hacker, C.E. Jorg: Ecological fitness, genomic islands, and bacterial pathogenicity. EMBO reports (2000)

[23] Bertelli, C., Gray, K.L., Woods, N., Lim, A.C., Tilley, K.E., Winsor, G.L., Hoad, G.R., Roudgar, A., Spencer, A., Peltier, J., Warren, D., Raphenya, A.R., McArthur, A.G., Brinkman, F.S.L.: Enabling genomic island prediction and comparison in multiple genomes to investigate bacterial evolution and outbreaks. Microb Genom 8(5) (2022) 10.1099/mgen.0.000818

[24] Bertelli, C., Laird, M.R., Williams, K.P., Simon Fraser University Research Computing, G., Lau, B.Y., Hoad, G., Winsor, G.L., Brinkman, F.S.L.: Islandviewer 4: expanded prediction of genomic islands for larger-scale datasets. Nucleic Acids Res 45(W1), 30–35 (2017) 10.1093/nar/gkx343

[25] Wood, D.E..S.L.S.: Kraken: ultrafast metagenomic sequence classification using exact alignments. BMC: Genome Biology 15 (2014)

[26] Bankevich, A., Nurk, S., Antipov, D., Gurevich, A.A., Dvorkin, M., Kulikov, A.S., Lesin, V.M., Nikolenko, S.I., Pham, S., Prjibelski, A.D., Pyshkin, A.V., Sirotkin, A.V., Vyahhi, N., Tesler, G., Alekseyev, M.A., Pevzner, P.A.: Spades: a new genome assembly algorithm and its applications to single-cell sequencing. J Comput Biol 19(5), 455–77 (2012) 10.1089/cmb.2012.0021

[27] Gardner, S.N., Slezak, T., Hall, B.G.: ksnp3.0: Snp detection and phylogenetic analysis of genomes without genome alignment or reference genome. Bioinformatics 31(17), 2877–8 (2015) 10.1093/bioinformatics/btv271

[28] Koren, S., Walenz, B.P., Berlin, K., Miller, J.R., Bergman, N.H., Phillippy, A.M.: Canu: scalable and accurate long-read assembly via adaptive k-mer weighting and repeat separation. Genome Res 27(5), 722–736 (2017) 10.1101/gr.215087.116

[29] Wick, R.R., Judd, L.M., Gorrie, C.L., Holt, K.E.: Unicycler: Resolving bacterial genome assemblies from short and long sequencing reads. PLoS Comput Biol 13(6), 1005595 (2017) 10.1371/journal.pcbi.1005595

[30] Darling, A.C., Mau, B., Blattner, F.R., Perna, N.T.: Mauve: multiple alignment of conserved genomic sequence with rearrangements. Genome Res 14(7), 1394–403 (2004) 10.1101/gr.2289704

[31] Hicks, J., Stuber, T., Lantz, K., Torchetti, M., Robbe-Austerman, S.: vsnp: a snp pipeline for the generation of transparent snp matrices and phylogenetic trees from whole genome sequencing data sets. BMC Genomics 25(1), 545 (2024) 10.1186/s12864-024-10437-5

[32] Walker, B.J., Abeel, T., Shea, T., Priest, M., Abouelliel, A., Sakthikumar, S., Cuomo, C.A., Zeng, Q., Wortman, J., Young, S.K., Earl, A.M.: Pilon: an integrated tool for comprehensive microbial variant detection and genome assembly improvement. PLoS One 9(11), 112963 (2014) 10.1371/journal.pone.0112963

[33] Tatusova, T., DiCuccio, M., Badretdin, A., Chetvernin, V., Nawrocki, E.P., Zaslavsky, L., Lomsadze, A., Pruitt, K.D., Borodovsky, M., Ostell, J.: Ncbi prokaryotic genome annotation pipeline. Nucleic Acids Res 44(14), 6614–24 (2016) 10.1093/nar/gkw569

[34] Tonkin-Hill, G., Lees, J.A., Bentley, S.D., Frost, S.D.W., Corander, J.: Fast hierarchical bayesian analysis of population structure. Nucleic Acids Research 47(11), 5539–5549 (2019) 10.1093/nar/gkz361

[35] Li, W., O’Neill, K.R., Haft, D.H., DiCuccio, M., Chetvernin, V., Badretdin, A., Coulouris, G., Chitsaz, F., Derbyshire, M.K., Durkin, A.S., Gonzales, N.R., Gwadz, M., Lanczycki, C.J., Song, J.S., Thanki, N., Wang, J., Yamashita, R.A., Yang, M., Zheng, C., Marchler-Bauer, A., Thibaud-Nissen, F.: Refseq: expanding the prokaryotic genome annotation pipeline reach with protein family model curation. Nucleic Acids Res 49(D1), 1020–1028 (2021) 10.1093/nar/gkaa1105

[36] Bertelli, C., Gray, K.L.e.a.: Enabling genomic island prediction and comparison in multiple genomes to investigate bacterial evolution and outbreaks. Microbial Genomics (2022) 10.1099/mgen.0.000818

[37] Wang, M., Liu, G., Liu, M., Tai, C., Deng, Z., Song, J., Ou, H.Y.: Iceberg 3.0: functional categorization and analysis of the integrative and conjugative elements in bacteria. Nucleic Acids Res 52(D1), 732–737 (2024) 10.1093/nar/gkad935

[38] Kuraku, S., Zmasek, C.M., Nishimura, O., Katoh, K.: aleaves facilitates on-demand exploration of metazoan gene family trees on mafft sequence alignment server with enhanced interactivity. Nucleic Acids Res 41(Web Server issue), 22–8 (2013) 10.1093/nar/gkt389

[39] Katoh, K., Rozewicki, J., Yamada, K.D.: Mafft online service: multiple sequence alignment, interactive sequence choice and visualization. Brief Bioinform 20(4), 1160–1166 (2019) 10.1093/bib/bbx108

[40] Li, H., Durbin, R.: Fast and accurate short read alignment with burrows-wheeler transform. Bioinformatics 25(14), 1754–60 (2009) 10.1093/bioinformatics/btp324

[41] Li, H., Handsaker, B., Wysoker, A., Fennell, T., Ruan, J., Homer, N., Marth, G., Abecasis, G., Durbin, R., Genome Project Data Processing, S.: The sequence alignment/map format and samtools. Bioinformatics 25(16), 2078–9 (2009) 10.1093/bioinformatics/btp352

[42] Jakob, S., Steinchen, W., Hanssmann, J., Rosum, J., Langenfeld, K., Osorio-Valeriano, M., Steube, N., Giammarinaro, P.I., Hochberg, G.K.A., Glatter, T., Bange, G., Diepold, A., Thanbichler, M.: The virulence regulator virb from shigella flexneri uses a ctp-dependent switch mechanism to activate gene expression. Nat Commun 15(1), 318 (2024) 10.1038/s41467-023-44509-z

[43] Kishida, K., Nonoyama, S., Lukas, T., Kawahara, S., Kudo, K., Nagata, Y., Ohtsubo, Y., Tsuda, M.: Conjugative transfer of incp-9 catabolic plasmids requires a previously uncharacterized gene, mpfk, whose homologs are conserved in various mpf(t)-type plasmids. Appl Environ Microbiol 85(24) (2019) 10.1128/AEM.01850-19

[44] Gonzalez-Montes, L., Del Campo, I., Garcillan-Barcia, M.P., Cruz, F., Moncalian, G.: Ardc, a ssdna-binding protein with a metalloprotease domain, overpasses the recipient hsdrms restriction system broadening conjugation host range. PLoS Genet 16(4), 1008750 (2020) 10.1371/journal.pgen.1008750

[45] Wen, A., Zhao, M., Jin, S., Lu, Y.Q., Feng, Y.: Structural basis of alpa-dependent transcription antitermination. Nucleic Acids Res 50(14), 8321–8330 (2022) 10.1093/nar/gkac608

[46] Lewis, J.A., Escalante-Semerena, J.C.: The fad-dependent tricarballylate dehy-drogenase (tcua) enzyme of salmonella enterica converts tricarballylate into cis-aconitate. J Bacteriol 188(15), 5479–86 (2006) 10.1128/JB.00514-06

[47] Hwang, S., Choe, D., Yoo, M., Cho, S., Kim, S.C., Cho, S., Choa, B.-K.: Peptide transporter csta imports pyruvate in escherichia coli k-12. Journal of Bacteriology 200 (2018) 10.1128/JB.00771-17

[48] Dix, S.R., Owen, H.J., Sun, R., Ahmad, A., Shastri, S., Spiewak, H.L., Mosby, D.J., Harris, M.J., Batters, S.L., Brooker, T.A., Tzokov, S.B., Sedelnikova, S.E., Baker, P.J., Bullough, P.A., Rice, D.W., Thomas, M.S.: Structural insights into the function of type vi secretion system tssa subunits. Nat Commun 9(1), 4765 (2018) 10.1038/s41467-018-07247-1

[49] Davies, J.S., Currie, M.J., Dobson, R.C.J., Horne, C.R., North, R.A.: Traps: the ‘elevator-with-an-operator’ mechanism. Trends Biochem Sci 49(2), 134–144 (2024) 10.1016/j.tibs.2023.11.006

[50] Rettner, R.E., Saier, J. M. H.: The autoinducer-2 exporter superfamily. J Mol Microbiol Biotechnol 18(4), 195–205 (2010) 10.1159/000316420

[51] Guedon, G., Libante, V., Coluzzi, C., Payot, S., Leblond-Bourget, N.: The obscure world of integrative and mobilizable elements, highly widespread elements that pirate bacterial conjugative systems. Genes (Basel) 8(11) (2017) 10.3390/genes8110337

[52] Zhang, D., Souza, R., Anantharaman, V., Iyer, L., Aravind, L.: Polymorphic toxin systems: Comprehensive characterization of trafficking modes, processing mechanisms of action, immunity and ecology using comparative genomics. Biology Direct 7(18) (2012)

[53] Iyer, L.M., Zhang, D., Rogozin, I.B., Aravind, L.: Evolution of the deaminase fold and multiple origins of eukaryotic editing and mutagenic nucleic acid deaminases from bacterial toxin systems. Nucleic Acids Res 39(22), 9473–97 (2011) 10.1093/nar/gkr691

[54] Ruhe, Z.C., Low, D.A., Hayes, C.S.: Polymorphic toxins and their immunity proteins: Diversity, evolution, and mechanisms of delivery. Annu Rev Microbiol 74, 497–520 (2020) 10.1146/annurev-micro-020518-115638

