## Supplementary figures and images for "Initial identification of genomic islands in *Taylorella equigenitalis* and *Taylorella asinigenitalis* and their distribution in isolates from around the world"

### Supplemental Figure

Supplemental Figure 1

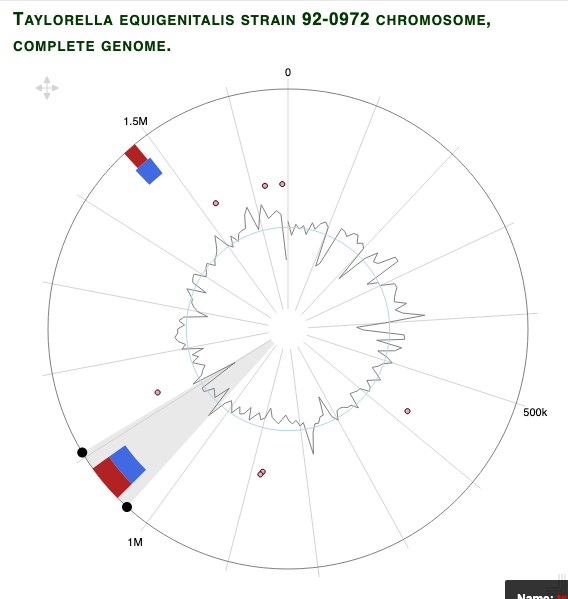

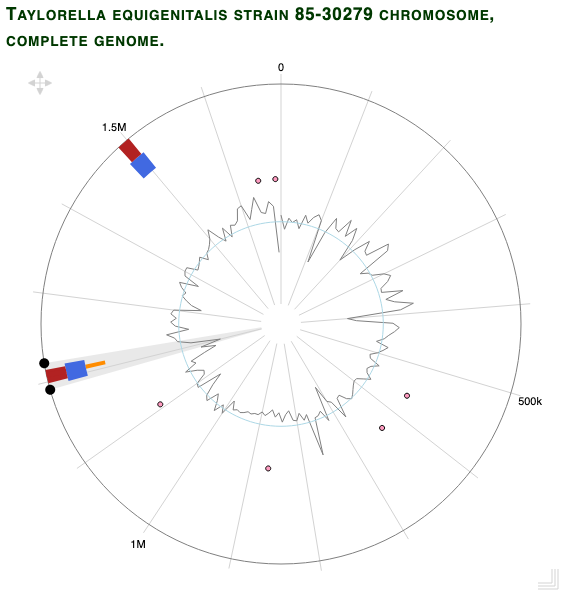

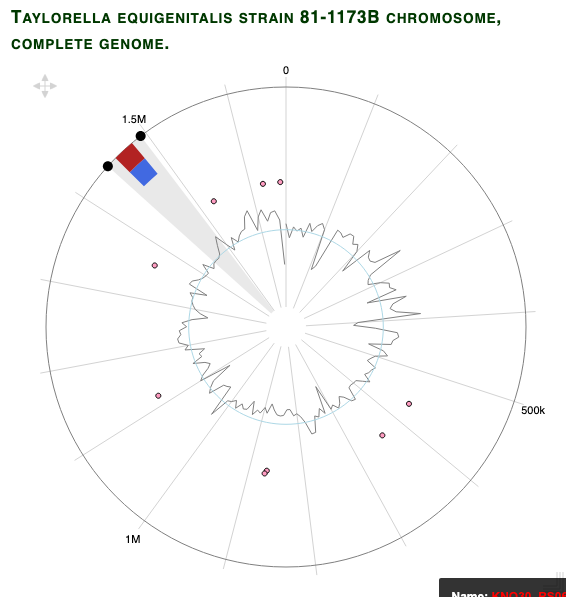

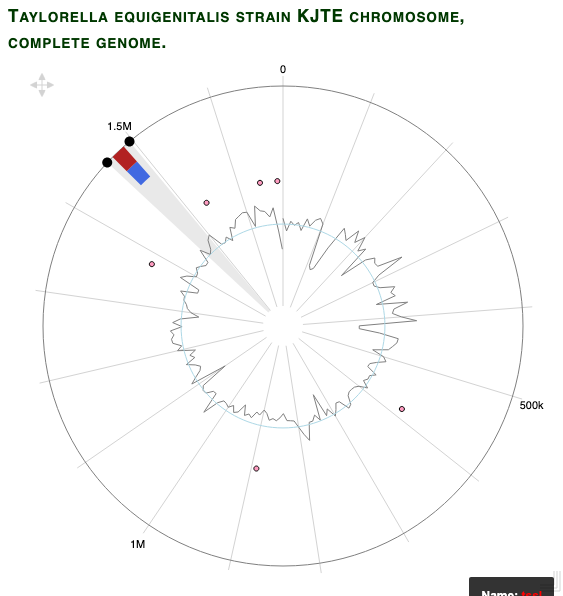

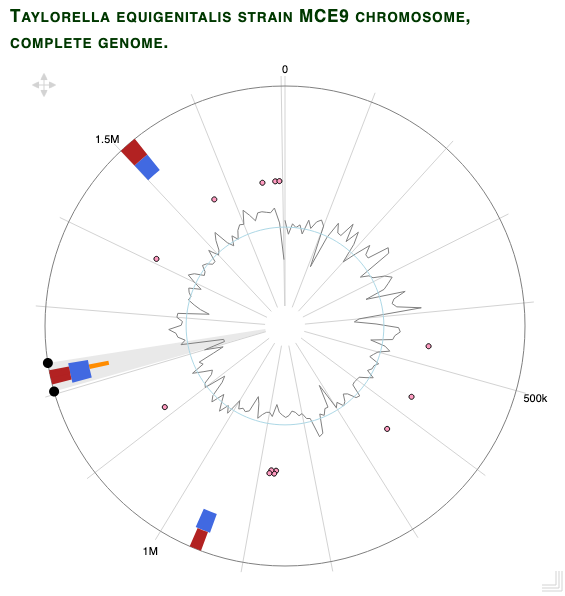

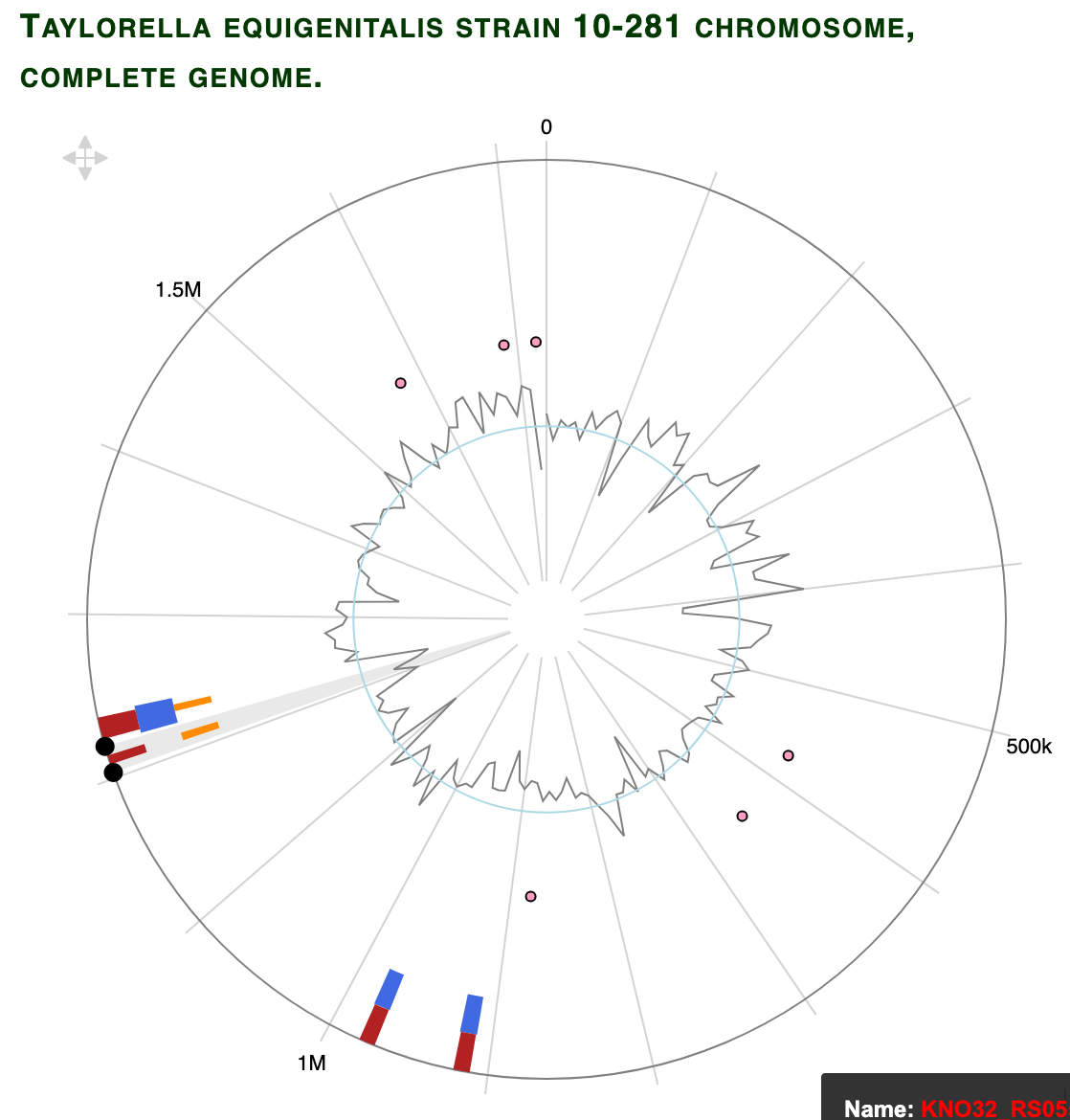

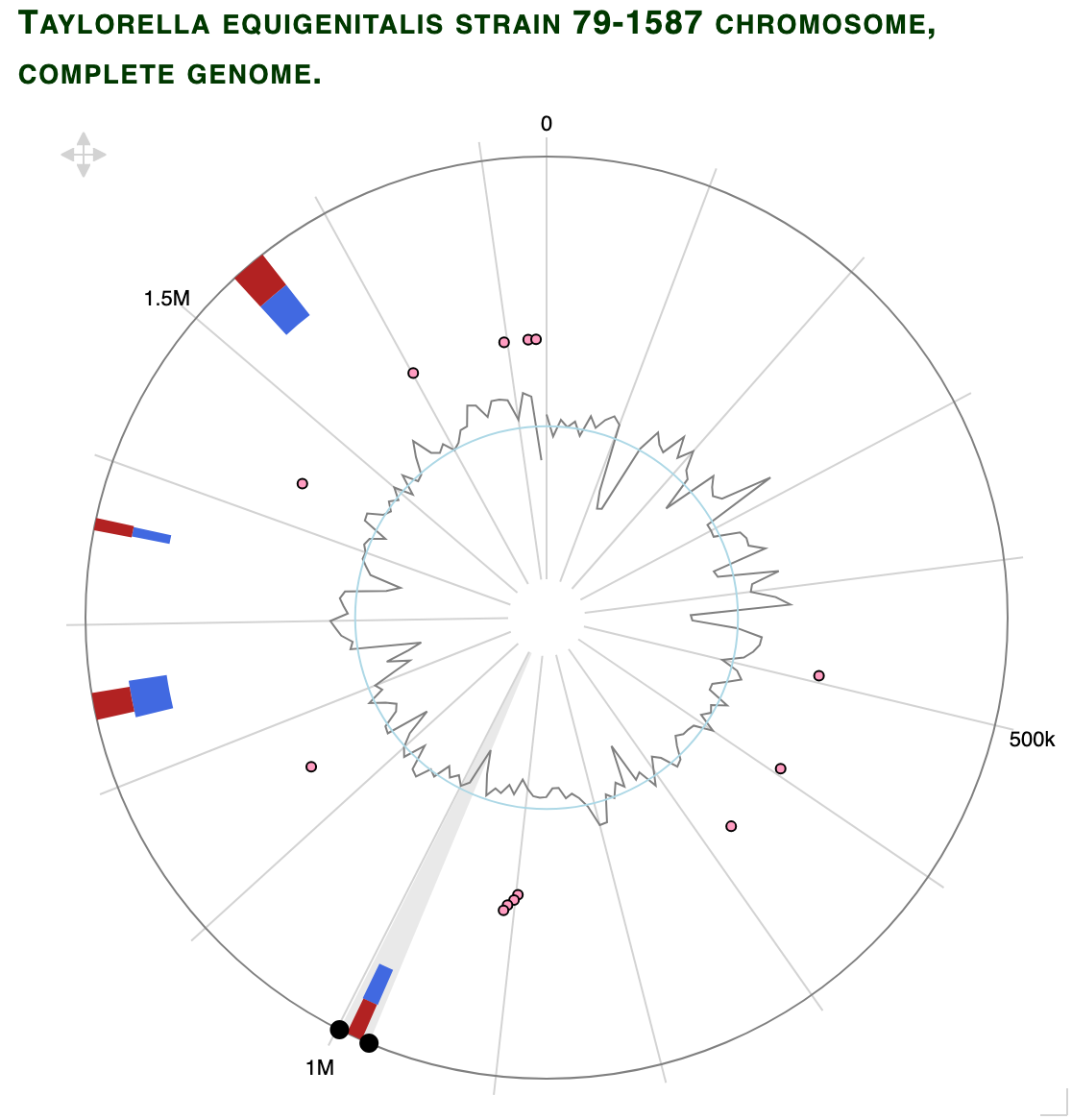

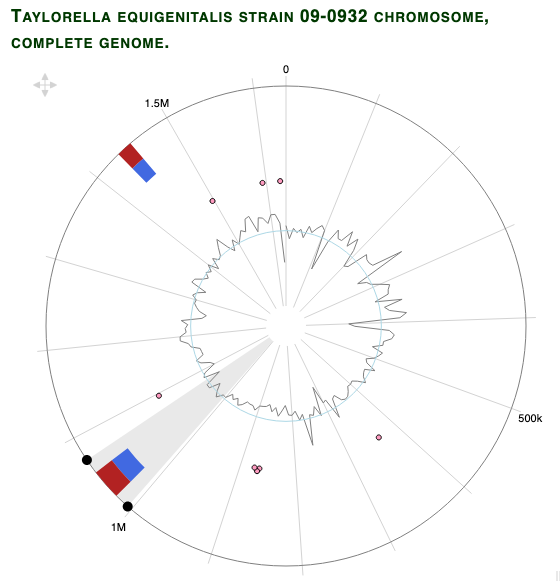

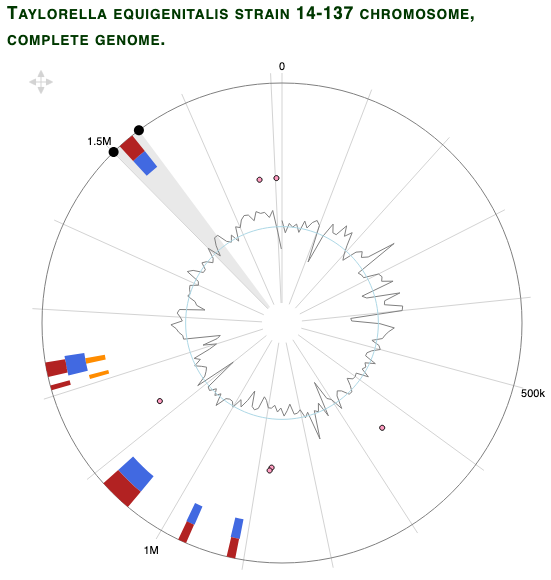

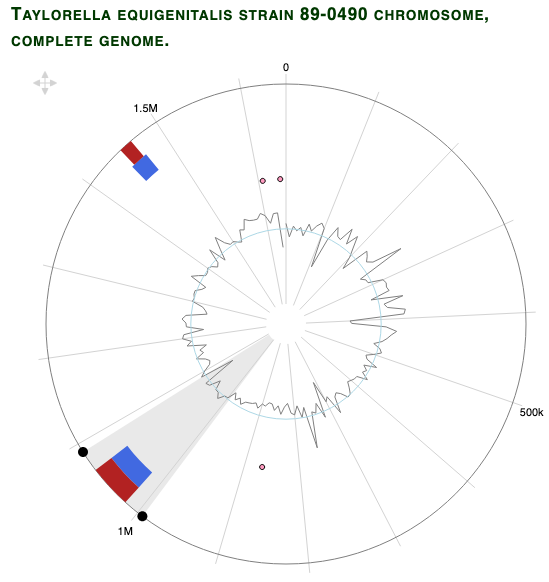

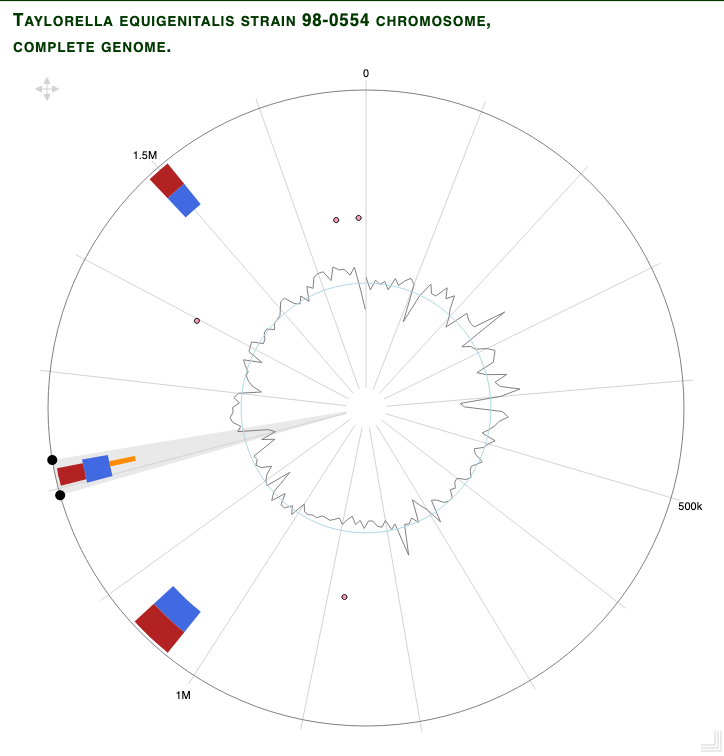

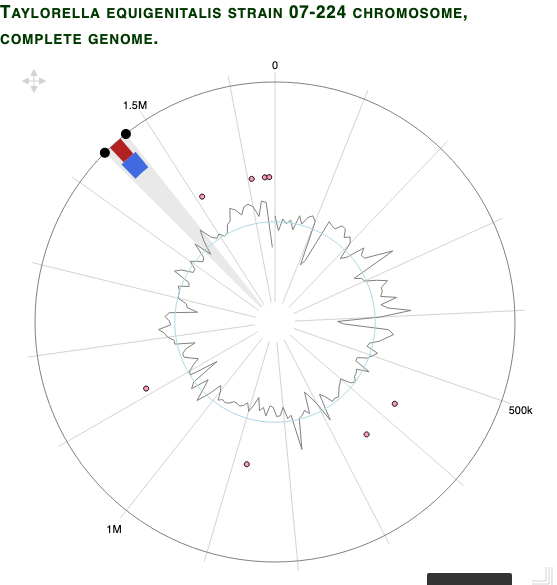
